# Accelerated Dynamic Time Warping on GPU for Selective Nanopore Sequencing

**DOI:** 10.1101/2023.03.05.531225

**Authors:** Harisankar Sadasivan, Daniel Stiffler, Ajay Tirumala, Johnny Israeli, Satish Narayanasamy

## Abstract

The design and supply of RT-PCR primers for accurate virus testing is a complex process. The MinION is a revolutionary portable nanopore DNA sequencer that may be used to sequence the whole genome of a target virus in a biological sample. Human samples have more than 99% of non-target host DNA and Read Until is a protocol that enables the MinION to selectively eject reads in real-time. However, the MinION does not have any in-built compute power to select non-target reads. SquiggleFilter is a prior work that identified the accuracy and throughput challenges in performing Read Until using the state-of-the-art solution and proposed a hardware-accelerated subsequence Dynamic Time Warping (sDTW) based programmable filter on an ASIC. However, SquiggleFilter does not work for genomes larger than 100Kb. We optimize SquiggleFilter’s sDTW algorithm onto the more commonly available GPUs. *DTWax* better uses tensor core pipes, 2X-SIMD FP16 computations and efficient data handling strategies using offline pre-processing, coalesced global memory loads, warp shuffles and shared memory buffering among other optimizations. *DTWax* enables Read Until and yields ~1.92X sequencing speedup and ~3.64X compute speedup: costup over a sequencing workflow that does not use Read Until.

## Introduction

With SARS-CoV-2 evolving and adapting to its new environment^1^ and becoming immune-evasive^2^, there is a possible threat from a variant that can evade our current gold standard tests and fuel a surge in cases. Reverse Transcription Polymerase Chain reaction (RT-PCR) is the current gold standard^3^ for SARS-CoV-2 diagnostic testing. Prior works have shown that RT-PCR requires the design and manufacture of custom PCR primers which is a complex, time-consuming, and error-prone process^4–6^. This limits the utility of RT-PCR in the early stages of a pandemic. Dunn and Sadasivan et al.^4^ developed SquiggleFilter, a portable virus detector that could be re-programmed to speed up the sequencing of reads from a viral target of interest. SquiggleFilter is an ASIC, envisioned to work alongside Oxford Nanopore Technology’s (ONT) MinION MK1B (or simply the MinION), a recent-to-market portable DNA sequencer that does not have any compute built into it. However, SquiggleFilter can only be programmed with references of size less than 100Kb and it being an ASIC, is not easily scalable.

Additionally, GPUs are becoming a more common choice for accelerated computing on sequencers– ONT sequencers GridION, PromethION, and MinION MK1C have GPUs built into them^7^. GPUs are also widely available in workplaces and on cloud platforms. While SquiggleFiter’s subsequence Dynamic Time Warping (sDTW) algorithm was optimized to work on an ASIC, we adapt and optimize it to work on the more common GPUs.

## Background

### Nanopore Sequencing

Long-read sequencing technology is increasingly becoming popular for rapid and accurate medical diagnosis^8, 9^ with lower adoption costs, end-to-end sequencing times, and improved portability and raw read accuracies. Long reads are particularly useful for applications that include structural variant calling and denovo assembly^8, 10^ as they, unlike short reads, can span highly repetitive regions in the genome.

Oxford Nanopore Technology’s (ONT) MinION is a long-read DNA sequencer that is low-cost, real-time, portable, and can perform digital target enrichment using software instead of time-consuming wet-lab based methods^4^. Just like any other sequencer, fragmentation of DNA (into reads) is an artifact of the wet-lab process. However, ONT sequencers can produce very long reads to help span the highly repetitive regions in the genome. Nanopore senses the DNA molecule that passes through the pore by measuring the characteristic disruptions in electric current density. Decoding this noisy but characteristic signal (squiggle) helps us understand the DNA base (A, G, T, or C).

MinlON is an ideal candidate for viral detection because of many factors^4^. MinION is portable, low-cost, and capable of real-time DNA sequencing. Unlike RT-PCR tests, where one has to perform enrichment of target DNA in low-concentration specimens in the wet-lab, MinION lets us save time and cost by digitally checking for targets while sequencing. In real-time, MinION can be controlled to selectively sequence just the target DNA strands and eject the non-target strands by reversing the electrical potential across the pore.

### Selective Sequencing

Most human samples have a very high fraction of non-target DNA (~99.9%. and most of it is human DNA)^11^. In order to save time and cost of sequencing, ONT has a feature called Selective Sequencing (Read-Until)^12^ which lets us selectively sequence only the target DNA reads while ejecting the non-target. As the read is sequenced, real-time compute may be performed to classify the read as a target or not. Non-targets are ejected by reversing the voltage in the nanopore while targets are completely sequenced.

The state-of-the-art selective sequencing pipeline^12, 13^ uses Guppy-fast for basecalling and Minimap2 for classifying the reads^13^ as shown in Fig. 1. Guppy is a deep neural network-based software that converts the raw signal output of the MinION (noisy squiggles) to bases. However, Guppy has a two-fold performance problem. Prior works have demonstrated how Guppy is slow and does not have the required throughput to handle the throughput of the MinION^4, 14–16^ and SquiggleFilter^4^ pointed out how the increasing throughput of the MinION amplifies this problem. We demonstrate the same problem in Fig. 2b. Secondly, Guppy is unable to accurately basecall small chunks of data. ~40% of the bases sequenced from a sample of average read length 2 Kbases is unclassified as shown in Fig. 2a because Guppy could not basecall these very short fragments accurately^16^.

**Figure 1.**
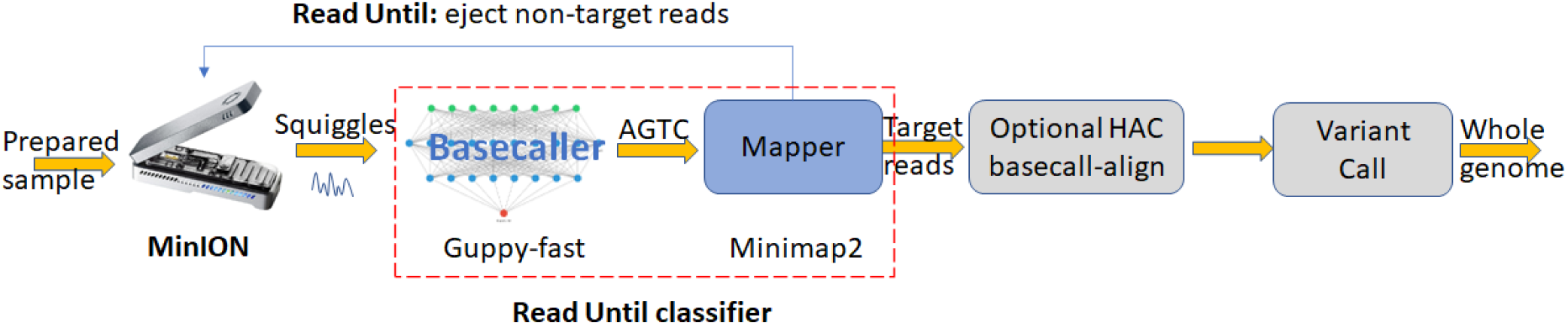
State-of-the-art selective sequencing pipeline uses Guppy-fast for basecalling and Minimap2 for classifying the reads.

**Figure 2.**
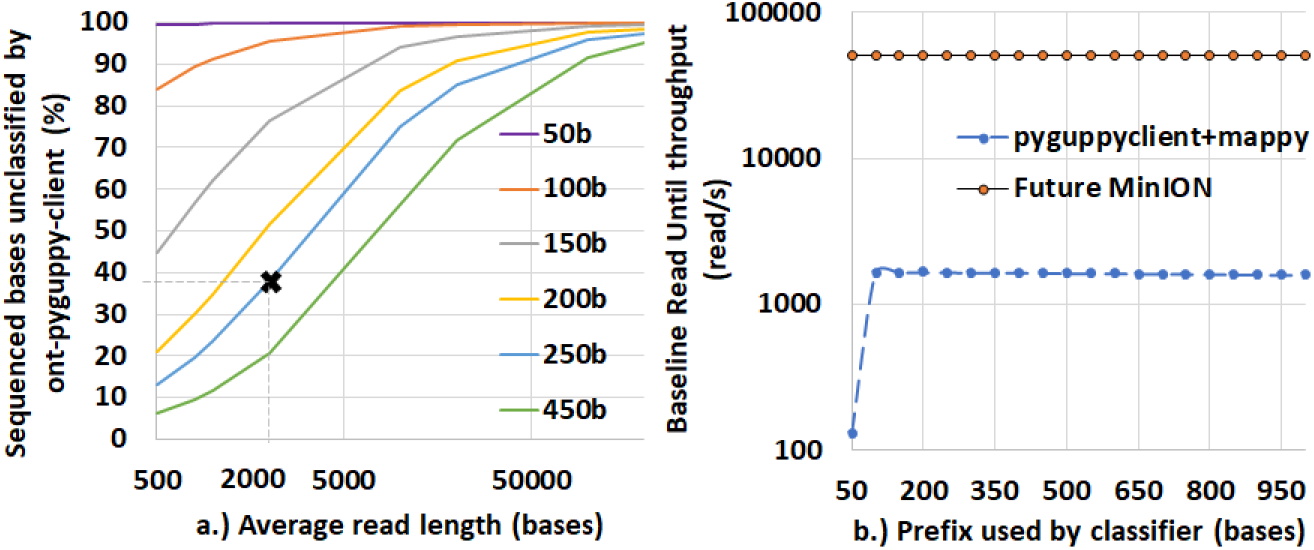
Guppy hascannot classify a high percent of sequenced bases and also has a throughput problem with Read Until. (a) ~40% of the bases sequenced are non-target in a 99.9: 0.1 non-target: target mix with an average read length of 2 Kbases. (b) Guppy followed by Minimap2 cannot match the throughput of a future MinION even on a high-end cloud instance which uses an A100 GPU for basecalling.

Hardware-accelerated SquiggleFilter^4^ was proposed as a replacement for the Read Until classification using Guppy followed by Minimap2.

### Prior Work

Since the MinION’s release in 2014, there has been a few attempts in improving the benefits of Read Until^4, 12–17^, of which only SquiggleFilter^4^ has the optimal combination of accuracy and throughput to keep up with a future MinION. The softwares performing Read Until can be broadly classified into two categories based on the inputs they operate on-signal space and basecalled space. Softwares operating in the signal space attempt to classify the input squiggle as a target or non-target while softwares operating in the basecalled space rely on the basecaller to transform raw squiggles to bases in real-time which is computationally expensive. We observe that the basecaller also basecalls smaller signal chunks poorly. The basecalled read prefixes may then be classified as a target or non-target.

Readfish^13^ and RUBRIC^12^ classify basecalled reads to detect targets using mapping tools like Minimap2. We show that ONT’s proprietary basecaller Guppy suffers from not being able to basecall ~10% of the reads confidently leading to them being unclassified by Minimap2. Additionally, Guppy, a deep neural network, also suffers from low throughput on high-end GPUs and cannot meet the real-time compute requirements of a future MinION.

Three of the signal space-based methods rely on event segmentation-a pre-processing step where raw squiggles are segmented into events to detect positions in the signal where we are more likely to see a new base. The very first attempt at Read Until^18^, UNCALLED^14^, and Sigmap^15^ use event segmentation as a pre-processing step. Loose et al.^18^ uses sDTW on python to perform Read Until from events on a CPU. This yields sub-optimal performance. UNCALLED follows up with FM-index look-ups and seed clustering to find a target map. Although UNCALLED has a good mapping accuracy for smaller genomes, We observe that UNCALLED does not have the necessary throughput to match the compute requirements of a future MinION. Sigmap does seeding followed by Minimap2-style chaining on the seeds to identify a target map. But we observe that Sigmap needs a relatively long read prefix to identify a sufficient number of seeds and this turns out to be 4000 raw samples. Additionally, we observe that Sigmap has lower mapping accuracy than UNCALLED.

SquiggleNet^19^ is a convolutional neural network-based software for classifying squiggles into target or non-target. However, SquiggleNet^19^ is slower than guppy followed by Minimap2 and only achieves similar mapping accuracy to Guppy followed by Minimap2. SquiggleFilter^4^ is a programmable ASIC that can match the throughput of a future MinION and yield optimal Read Until benefits. However, the initial cost of adoption is high as ASIC needs to be economically manufactured at scale and shipped in order to be deployed worldwide. GPUs on the other hand are already widely available at workplaces, shipped along with some of the sequencers, and also available on the cloud.

DTW has been parallelized in the past for various different applications on architectures including FPGAs^17, 20^, Intel Xeon Phis^21^, big data clusters^22^, customized fabrics^23^ and even GPUs^24, 25^. HARU^17^ is a recent work that implements SquiggleFilter’s algorithm on a budget-constrained FPGA. HARU cannot match the maximum throughput of the current MinION and is not a solution that can match the planned 100X throughput of the MinION. Crescent^26^ is a recent closed-source implementation of SquiggleFilter’s algorithm directly on the GPU but ends up being 29.5X lower in throughput than *DTWax* possibly because of several reasons including not utilizing warp synchronized register shuffles for data sharing between threads and fewer number of cells computed per thread. cuDTW++^27^ is the best-performing prior work on GPU which accelerates DTW. However, cuDTW++ is ~2.6X slower than DTWax and is built for database querying of very small queries and not for subsequence Dynamic Time Warping that is required to perform Read Until. Additionally, the normalization step is performed very inefficiently.

### subsequence Dynamic Time Warping

sDTW is a two-dimensional dynamic programming algorithm tasked with finding the best map of the whole of the input query squiggle in the longer target reference. In sDTW’s output matrix computation, parallelism exists along the off-diagonal of the matrix and therefore, the computation happens in a wavefront parallel manner along this off-diagonal. Diagonals are processed one after the other. If the query is assumed to be along the vertical dimension of the matrix and the target reference along the horizontal dimension, the minimum score on the last row of the matrix will point to the best possible map of the query to the reference. This score may be compared to a threshold to figure out if the query is a target or not. The sDTW cost function is defined as follows:

### SquiggleFilter

While traditional RT-PCR tests rely on complex custom primer design and time-consuming wet-lab processes for target enrichment, MinION can be controlled to selectively sequence only the target virus of interest using the Read Until feature. Utilizing the Read Until feature requires making real-time classifications during sequencing but the current MinION does not have any compute power. Dunn and Sadasivan et al.^4^ demonstrated how basecalling is the bottleneck and constitutes ~88-96%. of Read Until assembly and how this problem was amplified with the projected 100X increase in ONT’s sequencing throughput. Their solution, SquiggleFilter^4^, uses hardware accelerated subsequence Dynamic Time Warping (sDTW) to perform Read Until.

SquiggleFilter addresses the compute bottlenecks in portable virus detection and is designed to even handle the higher throughput of a future MinION. SquiggleFilter is programmable and offers better pandemic preparedness apart from saving time and cost of sequencing and compute. However, SquiggleFilter’s limited on-chip memory buffer only lets it test for viral genomes smaller than 100Kb. Additionally, SquiggleFilter uses a modified version of sDTW algorithm where the accuracy dip from various hardware-efficiency focussed optimizations are overcome with the match bonus^4^. Match bonus is a solution to a problem on the ASIC and performing this on the GPU can introduce branch divergence. We eliminate the match bonus, retain the assumption of reference deletions and optimize sDTW to run on GPUs.

### Our contributions

In this work. we present *DTWax,* a GPU-accelerated sDTW software for nanopore Read Until to save time and cost of nanopore sequencing and compute. We adapt SquiggleFilter ASIC^4^’s underlying sDTW algorithm to suit a GPU in order to overcome the limitation with reference lengths SquiggleFilter had. While sDTW was optimized for integer compute on the ASIC, we fine-tune sDTW for high throughput on the GPU. While SquiggleFilter uses integer arithmetic and Manhattan distances on the ASIC, we use floating point operations and Fused-Multiply-Add operations on the GPU. We also demonstrate how to utilize some of the GPU’s high throughput tensor core’s compute power for non-ML workloads.

As a first step, we speed up the online pre-processing step (normalization) on FP32 tensor cores using the batch normalization functionality from the CUDNN library traditionally used for machine learning workloads. *DTWax* is optimized to make use of the high throughput Fused-Multiply-Add instructions on the GPU. Further, we use FP16 and FP16 tensor core’s Matrix-Multiply-Accumulate (MMA) pipe for higher throughput for sDTW calculation. Using FP16 helps us process the forward and reverse strands, thereby extracting more parallelism to help improve the latency and throughput of classification. We also make use of offline pre-processing of reference squiggle index for coalesced loads, cudastreams for better GPU occupancy, intra-, and inter-read parallelism, register shuffles, and shared memory for low-overhead communication while processing the same query.

*DTWax* achieves ~1.92X sequencing speedup and ~3.64X compute speedup: costup from using nanopore Read Until for a future MinION (with 100X the current throughput) on an A100 compared to a workflow that does not use Read Until.

## Methods

### Offline pre-processing

ONT has published a k-mer current model^28^ which provides a reference to map a 6-mer to an expected value of the current output from the MinION. We use this k-mer model to map the reference genome of the target virus to a noise-free FP16 squiggle reference. The squiggle reference will be of length (target_length −6+1). We also pack two FP16 values (one from the forward strand and another from the reverse strand) into a __half2 reference word (built-in CUDA datatype of two FP16 half-words). Further, we make use of the prior knowledge of the target reference to ensure coalesced global memory reads by re-ordering the target reference offline.

### Online pre-processing: Normalization

The output squiggle of the MinION (query) is read from ONT’s proprietary FAST5 file format. The raw integer data is then scaled to pico-amperes (float32). The first few samples (1000) are trimmed to cut adapters and barcodes off. We re-purpose the CUDNN-Batchnorm to z-score normalize the 1-dimensional FP32 query current signal. CUDNN utilizes tensor cores and performs normalization at a very high throughout (~6X the throughput of sDTW). The signal is then rounded off to FP16 and copied into a __half2.

### DTWax: architecture

We adopt the segmented-sDTW architecture introduced in prior works^27^ where each segment is a fixed number of cells in a row whose scores are computed by a thread. DTWax breaks down the processing of longer target references into multiple sub-matrices, each processing a fixed number of target bases. The reference length processed per sub-matrix is configurable and is set to 832 bases for optimal performance on an A100. Within a sub-matrix, each thread is responsible for processing a configurable but equal number of cells (cells per thread is called a segment). Wavefront parallelism exists along the off-diagonal segments in the sub-matrix as shown in Fig.3. Thread 0 is the first to finish its computation inside the sub-matrix while thread 31 is the last to finish. Target reference is loaded into registers (one FP16 reference sample each for forward and reverse strands into a single __half2 datatype) from global memory using coalesced loads.

**Figure 3.**
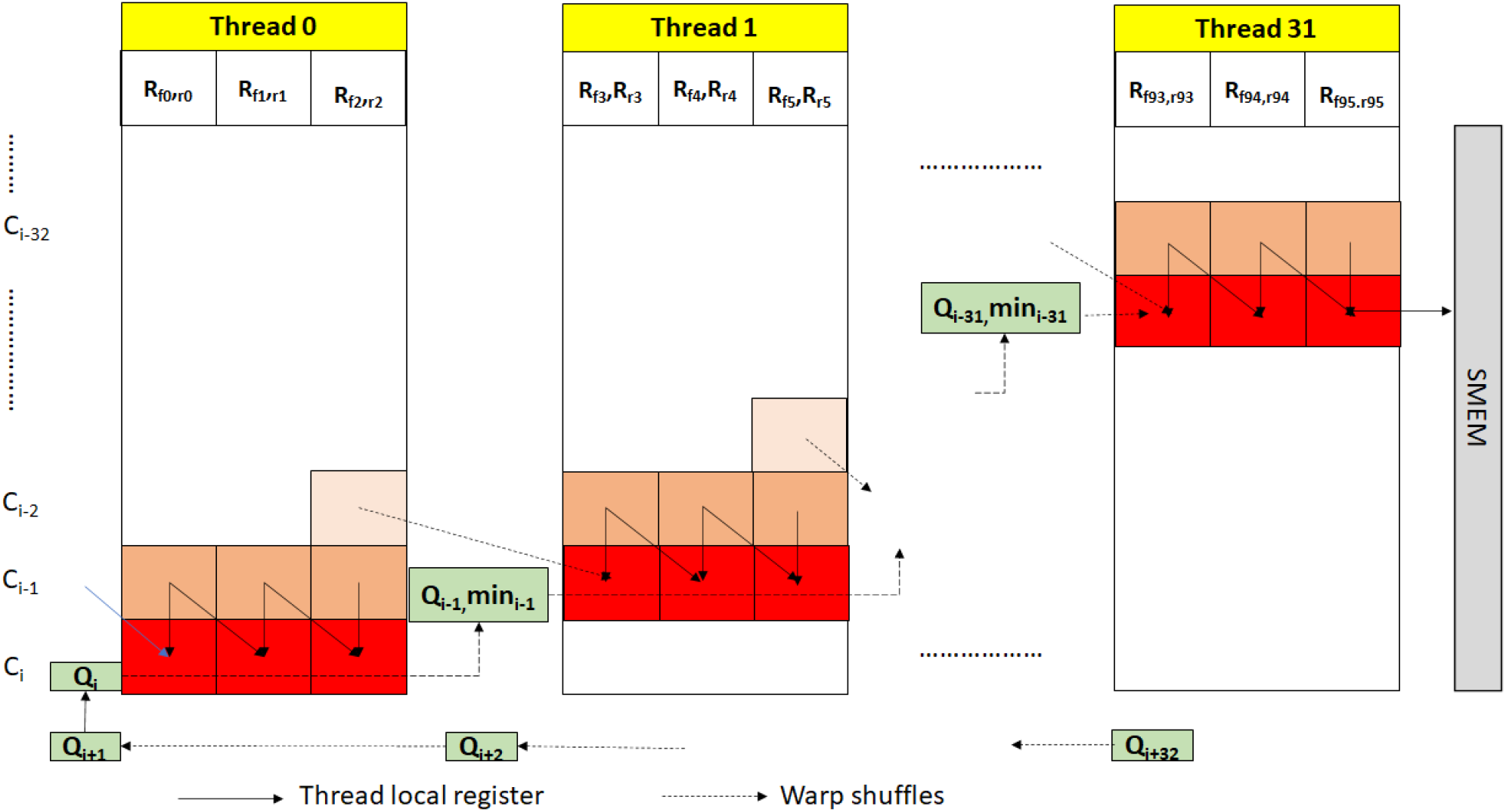
Efficient intra- and inter-matrix communication in *DTWax*.

For intra-sub-matrix communication, we exploit warp shuffles for efficient register-to-register transfers within the same warp. This is an idea demonstrated by Schmidt et al^27^ but not completely explored. Threads in a warp use warp shuffles to transfer the query sample, the minimum score of the segment, and the score of the last cell in the segment to the thread on its right. Instead of using a global reduction to find the final minimum score for *DTWax,* we use the efficient warp-shuffles to pass the minimum scores of the segments between threads. Inter-sub-matrix communication happens via shared memory transfers instead of relying on global memory. A thread block processing a read writes the last column of the sub-matrix into the shared memory and reads it back while calculating the consecutive sub-matrix for the same read.

### Intra- and inter-read parallelism

Using all the warps on an SM to process a single query would mean that the last warp remains idle and is ineligible for compute for an initial period of time. Therefore, we choose to process one read with a thread-block of only 32 threads. We have intra- and inter-read parallelism. Every query is processed by a thread block of 32 threads. Within a thread block, we have intra-read parallelism from 32 parallel threads each computing a segment of the sub-matrix. Across the GPU, we have inter-read parallelism as there as multiple concurrent blocks operating on different reads on any given Streaming Multi-processor (SM).

### Coalesced global memory access

The offline re-ordering of the target reference enables us to perform coalesced reads from global memory (as many loads as the length of one segment in a sub-matrix) before computation starts in the sub-matrix. The normalized target reference is an array of __half2 datatype. This enables the vectorized processing of the input query signals on the high throughput FP16 pipe. Additionally, after the normalized query is read from the global memory in chunks of 32 __half2 query samples using a coalesced load of 128B, it is then efficiently transferred between threads of a warp using warp shuffles.

### FP16 for 2X throughput

SquiggleFilter^4^ has demonstrated that the information from the ONT sequencer may be captured using 8 bits. While the ASIC was custom-designed for integer arithmetic, GPUs are designed for high throughput floating point arithmetic. Among the floating point pipes available, we use the high throughput FP16 pipe on A100 (2X throughput compared to FP32) for *DTWax.* Computation with respect to the forward strand of the target reference happens on the first FP16 lane while the second FP16 lane computes with respect to the reverse strand. Utilizing __half2 FP16 pipes (FP16 vectorization) not only helps us to increase throughput but also improves the latency by 2X because we concurrently process both the forward and the reverse strand of the target with respect to the query in every cell of the sub-matrix.

### Utilizing tensor core pipe

HFMA2.MMA pipe on the tensor core has one of the highest throughputs on A100. We re-formulate the addition in the cost function of *DTWax* to a Fused Multiply-Add operation in order to utilize the otherwise under-utilized HFMA2.MMA pipe. We are then able to better throttle the compute instructions between HFMA2.MMA and the remaining FP16 pipes instead of increasing the traffic on the FP16 pipe.

### Assuming no reference deletion

Using the same assumption from SquiggeFilter^4^ that viral strains have minimal reference deletions, we observe our accuracy of mapping using *DTWax* improves and our new cost function becomes simpler as we now only have to find a single minimum per cell instead of two minimums. Line 7 from Algorithm 1 is simplified to:

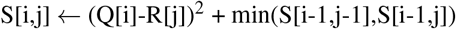

#### Algorithm 1 sDTW algorithm

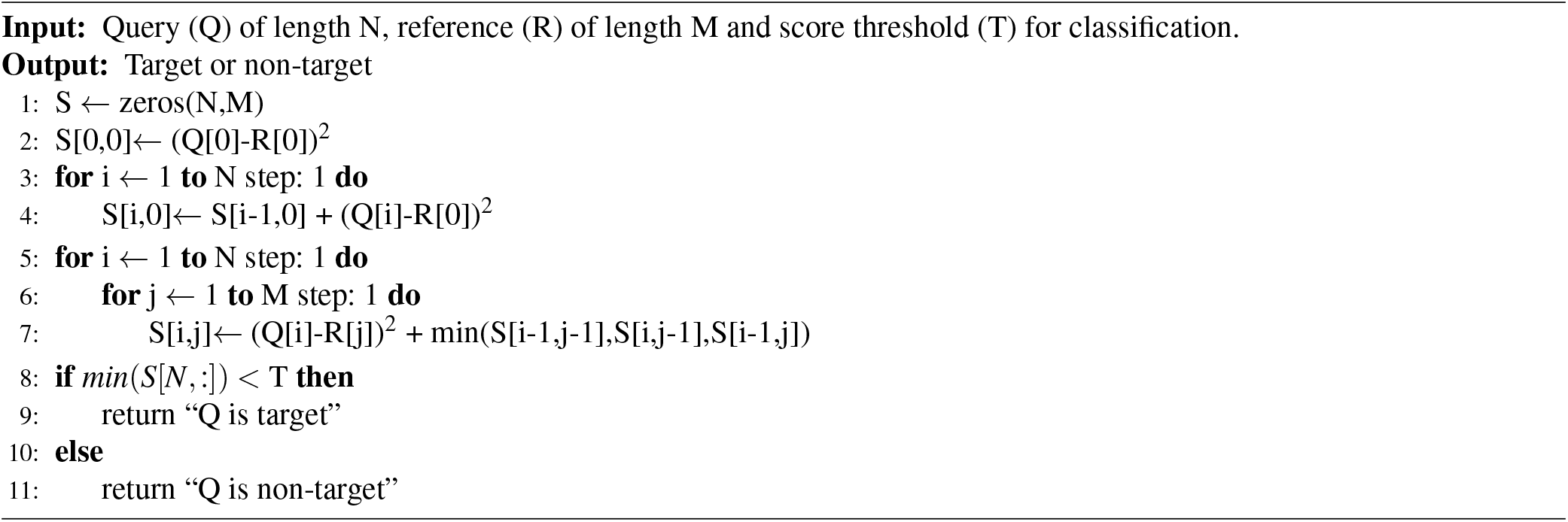

### Optimizing occupancy and branch divergence

We ensured high SM utilization by finding the right balance between the number of resident warps on the SM and shared memory utilization. Further, we keep the GPU occupancy high by issuing concurrent asynchronous workloads to the GPU using cudastreams. Memory transfers to and from the GPU are overlapped with compute on the GPU.

We reduce the branch divergence via partial loop unrolling. The first sub-matrix does not read from shared memory and the last sub-matrix does not write into shared memory. Unrolling the first and last sub-matrix computations of the query-target matrix helps improve performance.

### Configurability and scalability

*DTWax* can be reprogrammed to test for any target reference of interest. Unlike some of the prior works^4, 27^, *DTWax* can be reporgrammed to test for longer target references. Further, one may easily try and scale *DTWax* across multiple GPUs for higher throughput on longer or multiple target references.

## Implementation

### Experimental Setup

For all our GPU evaluations we use an NVIDIA A100 GPU on Google Cloud Platform’s (GCP) a2-highgpu-1g instance with 40GB of memory and 85GB of host memory. Our CPU baselines are evaluated with hyper-threading enabled on GCP c2-standard-60 (30 Intel Cascade Lake cores and 240 GB of memory). pyguppy-client and *DTWax* use GPU while other software use CPU.

NVIDIA Nsight Systems^29^ is used to visualize concurrent CUDA events, and NVIDIA Nsight Compute^30^ is used to profile GPU events.

We use public datasets used by SquiggleFilter^4^. We sequenced lambda phage DNA on a MinION R9.4.1 flow cell following the Lambda Control protocol at the University of Michigan’s laboratory using the ONT Rapid Library Preparation Kit^31^. Human datasets (sequenced with MinION R9.4 and R9.4.1 flow cells) are obtained from ONT Open datasets^32^ and the Nanopore Whole-Genome Sequencing Consortium^33^.

We use UNCALLED v2.2, Sigmap v0.1, Minimap2 v2.17, pyguppy-client v0.1.0 and Centrifuge v1.0.4-beta. We optimally configure the software for better mapping accuracy. We configure Sigmap for better event detection by setting “–min-num-anchors-output 2 –step-size 1”. We turn off minimizers in Minimap2 for better mapping accuracy by setting “-w=1 -k=15”.

### Optimal GPU configurations

In order to optimize *DTWax*’s throughput on A100, we introduce more inter-read parallelism by fitting 32 blocks on each of the 108 Streaming Multiprocessors on the GPU. We process one read per thread block of size 32 threads. We observe that having multiple concurrent warps per read resident on the SM may not be beneficial as there may be a long latency in the last warp of a read getting valid inputs from the previous warp and starting to calculate useful values. Using Nsight Compute, we maximize the number of reference bases processed per thread (segment size) to 26 amounting to a total of 32×26 reference bases processed per thread block. In order to reduce global memory transactions, we use shared memory for inter-sub-matrix communication. A shared memory of size 500B is allocated per block to store the output of the last column of a sub-matrix calculated by the last thread in a thread block. This is read by the thread block when it processes the next set of 32×26 reference bases.

### Incremental Optimizations

*DTWax* is a compute-bound kernel. Fig.4 gives the reader a better understanding of the relative benefits of some of the main optimizations that go into *DTWax*.

**Figure 4.**
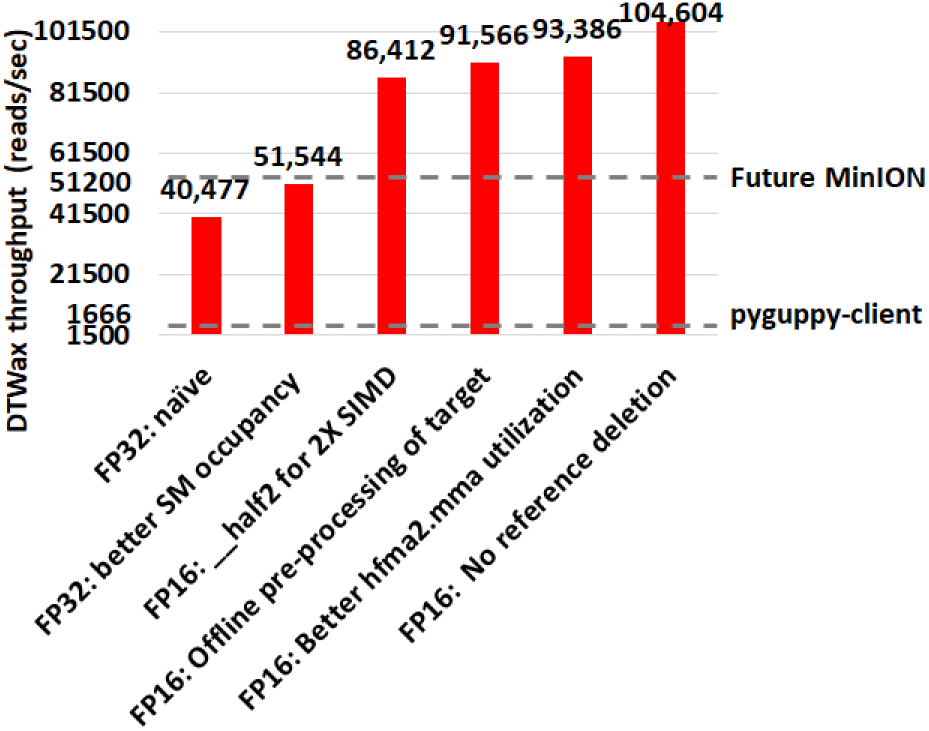
A series of performance optimizations enables *DTWax* to handle more than 2X the projected future sequencing throughput.

## Results

We use two metrics to evaluate the benefits of using *DTWax*–sequencing speedup and compute speedup: costup. Sequencing speedup is defined as the speedup in the end-to-end sequencing time from using *DTWax* for Read Until over a conventional nanopore sequencing workflow that does not use Read Until. Accessing an NVIDIA A100 GPU instance on the cloud is priced ~10% higher than the CPU instance we use for benchmarking. Hence, we normalize the compute time savings from using *DTWax* for Read Until to the cost of the cloud GPU instance to estimate compute speedup: costup. Further, we also compare the F1-score of *DTWax* in making Read Until classifications.

*DTWax* yields up to ~1.92X sequencing speedup and ~3.64X compute speedup: costup with a future MinION of 100X throughput when compared to a sequencing workflow that does not use Read Until as shown in Fig. 5. Additionally, we observe that using a prefix length of 250 bases yields the best benefits from using Read Until on a dataset of average read length 2 Kbases. One may also observe that savings from ont-pyguppy client degrades with increasing read-prefix lengths used for classification because ont-pyguppy cannot process streaming inputs and concatenate to outputs. Therefore, ont-pyguppy-client has to basecall the entire prefix again. Sigmap is unable to extract events from signals less than 400 bases. Please note that *DTWax* and pyguppy-client are GPU solutions and we label them using dotted lines on all the plots.

**Figure 5.**
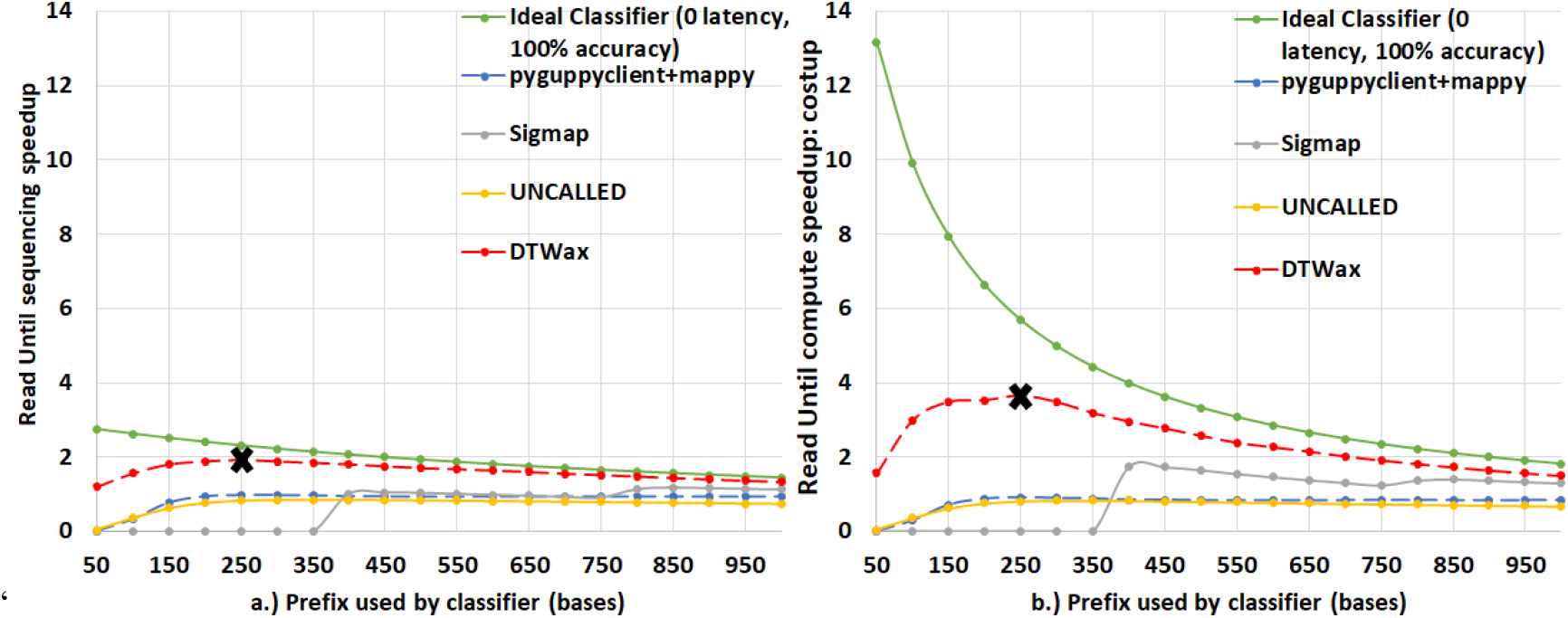
*DTWax* yields ~1.92X sequencing speedup and ~3.64X compute speedup: costup. (a) *DTWax* yields the best sequencing speedup of ~1.92X over a conventional pipeline that does not use Read Until on reads of length 2000 bases. (b)DTWax yields the best compute speedup: costup of ~3.64X. Prefix length used for classification is 250 bases for the best performance in both cases.

Fig. 6 shows that *DTWax* can handle more than 2X the throughput of a future MinION and has ~7.18X lower latency than pyguppy-client followed by mappy using a prefix length of 50 bases. Because of this, we operate *DTWax* at a processing granularity of 50 bases over a prefix length of 250 bases to make a Read Until classification decision. To explain the performance of pyguppy-client on longer prefixes, we profile pyguppy-client using NVIDIA nsight-compute and observe that the GPU occupancy is higher with longer prefix lengths resulting in better benefits from longer prefix lengths. UNCALLED seems to have a very high one-time fixed cost for path buffer management for storing output forest of trees and there is negligible cost with added prefix lengths. Although, Sigmap is the best in terms of throughput and latency, it cannot extract useful information from 250 bases and has lower F1 scores in classifying prefix lengths longer than 400 bases as shown in Fig. 7.

**Figure 6.**
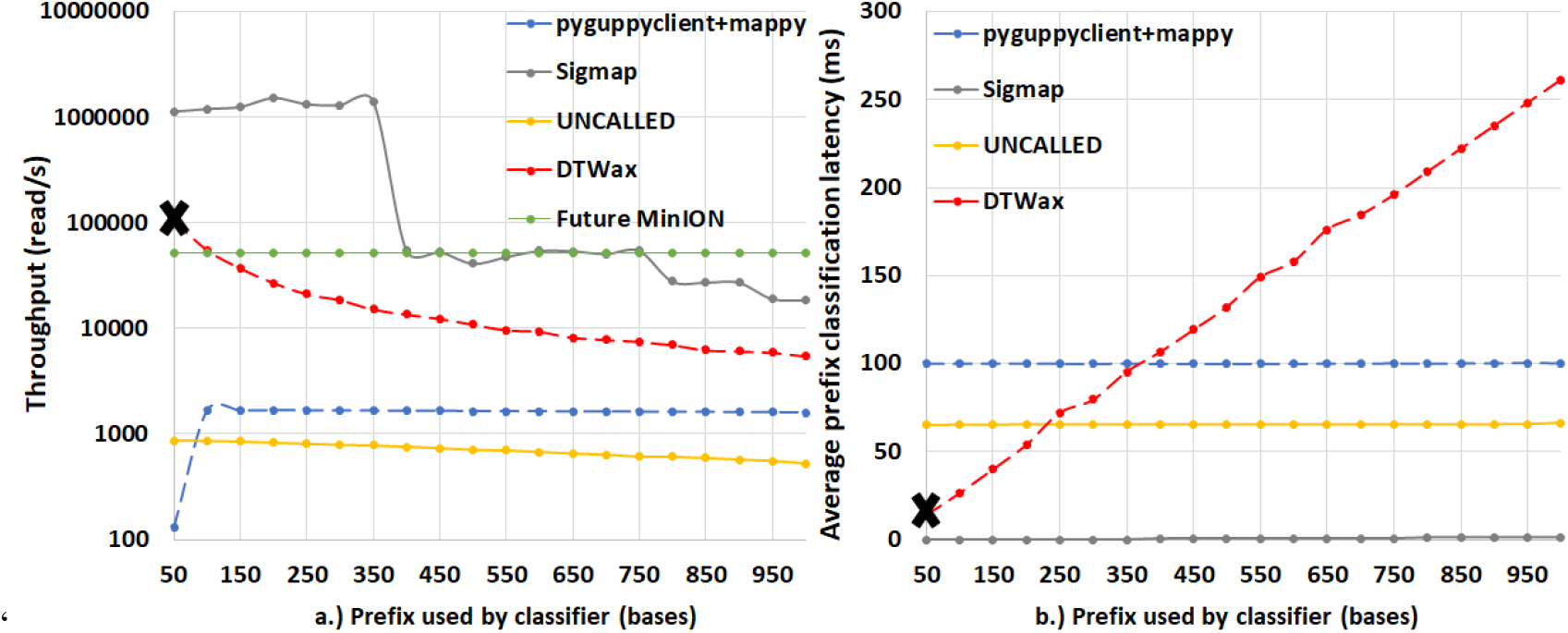
*DTWax* can handle more than 2X the throughput of a future MinION and has ~7.18X lower latency than pyguppy-client followed by mappy. (a)Unlike pyguppy-client followed by mappy, *DTWax* operating at a granularity of 50 bases can handle twice the throughput of a future MinION (b)*DTWax* takes only ~14 milliseconds to classify 50 bases and is ~7.18X faster than pyguppy-client followed by mappy.

**Figure 7.**
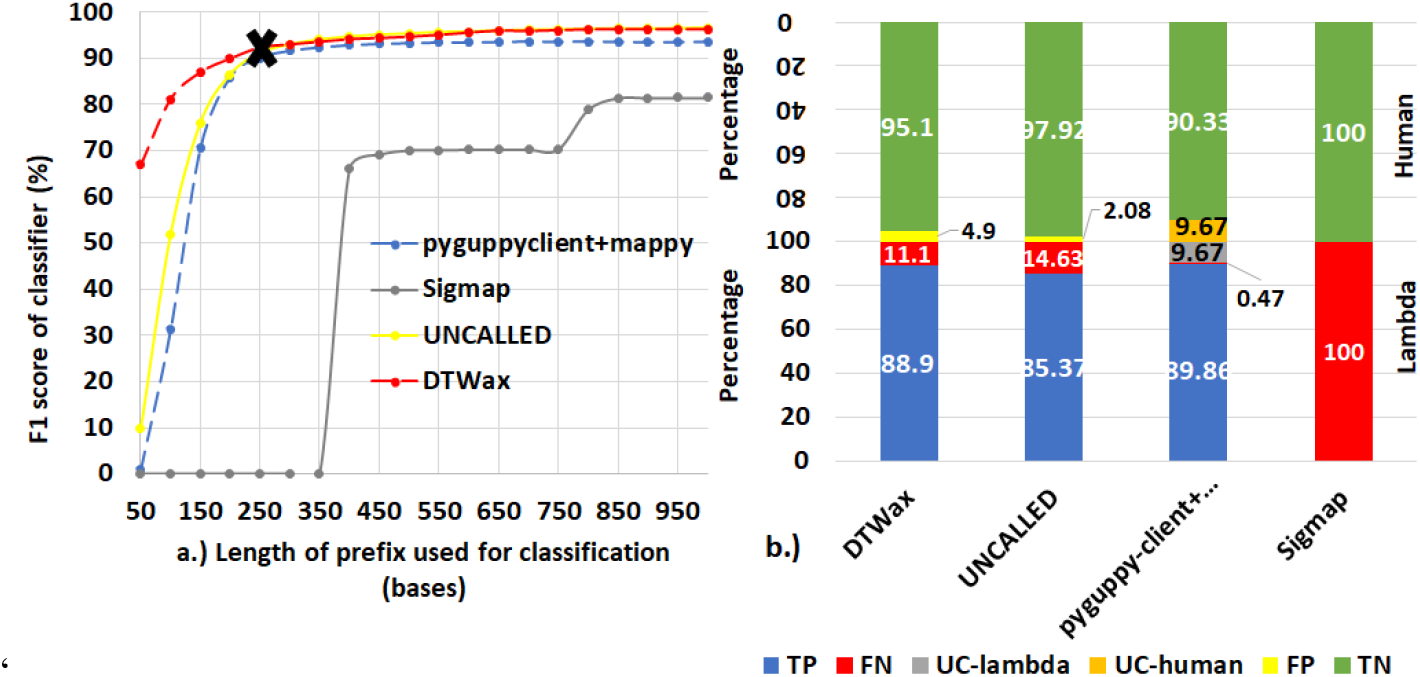
*DTWax* is the most accurate Read Until classifier. (a) *DTWax* has an F1-score of ~92.24% and is the most accurate Read Until classifier using a prefix length of 250 bases. (b) *DTWax* is better at filtering non-target (human) reads out than pyguppy-client followed by mappy while being almost comparable in succesfully retaining target reads

*DTWax* is the most accurate Read Until classifier as shown in Fig. 7. The sequencing speedup and compute time savings from *DTWax* are higher for longer reads as shown in Fig. 8.

**Figure 8.**
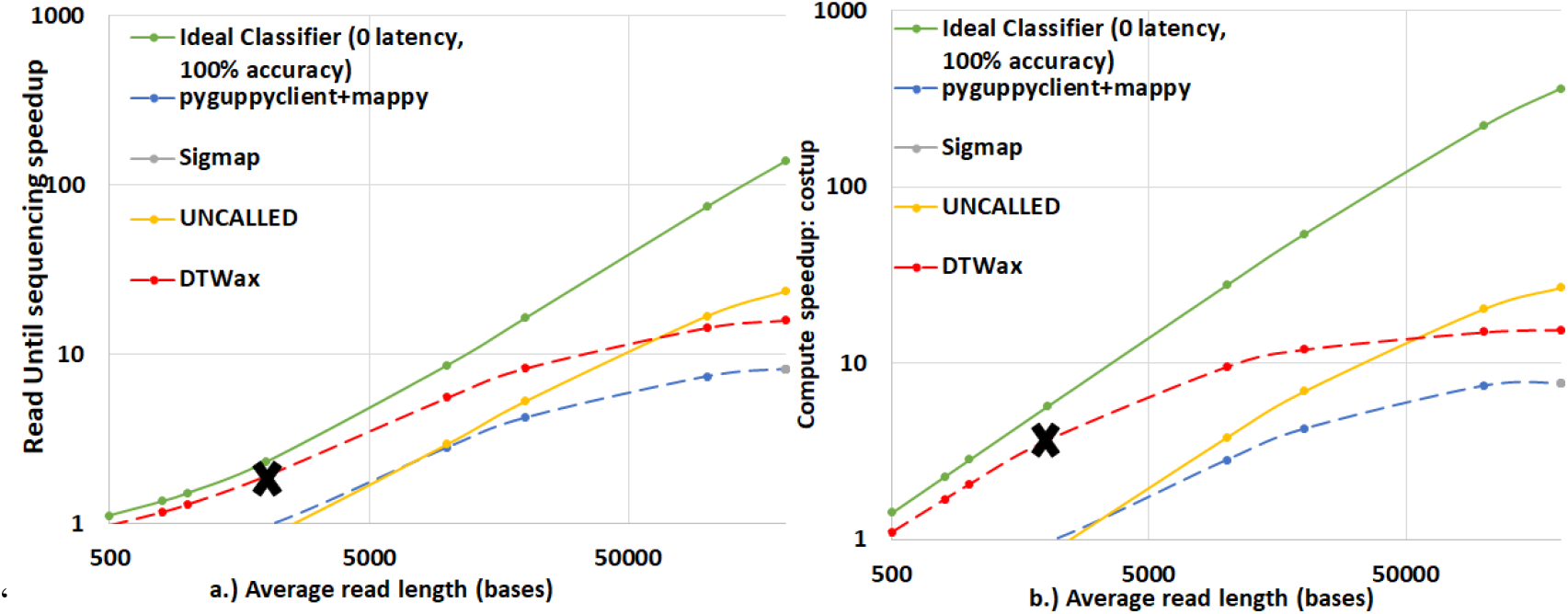
*DTWax* yields higher benefits on longer read lengths. In (a) and (b), we see increasing benefits from using Read Until on higher read lengths and *DTWax* is the best solution for read lengths shorter than ~50Kbases.

## Conclusions

SquiggleFilter is an ASIC that can be programmed to perform pathogen detection with the portable MinION sequencer. However, the ASIC cannot be programmed with references longer than 100 Kb. We adapt SquiggleFilter’s underlying sDTW algorithm to the more widely available GPUs to democratize Read Until. To make sDTW performant on the GPU, we do offline pre-processing of target reference to ensure coalesced loads from global memory, reduce branch divergence, and utilize FP16 vectorization and tensor core pipes. Further, we use warp-shuffles for efficient intra-sub-matrix communication and shared memory for low-latency inter-sub-matrix communication. Further, we assume no reference deletions to improve both the throughput and F1-score. We show that *DTWax* on an NVIDIA A100 GPU achieves ~1.92X sequencing speedup and ~3.64X compute speedup: costup over a sequencing workflow that does not use Read Until.

## Acknowledgements

Development of *DTWax* was supported by NVIDIA Corporation and the University of Michigan Ann Arbor (via D. Dan and Betty Kahn foundation grants). Additionally, we would like to thank Google Cloud Platform for awarding us cloud research credits for the final evaluations of this research.

## Author contributions statement

H.S. performed the analysis, design, implementation and evaluation of *DTWax* software apart from writing the manuscript. D.S. and A.T. recommended various performance optimizations and CUDA best practices for implementing *DTWax* based on DTWax’s profile. J.I. and S.N. led the collaborative effort and helped design the optimization strategy for *DTWax.* All authors reviewed the manuscript.

## Data availability statement

Datasets used are published by SquiggleFilter here::https://doi.org/10.5281/zenodo.5150973. ONe may download the dataset using the scrip provided from SquiggleFilter: https://github.com/TimD1/SquiggleFilter/blob/master/setup.sh

## Additional information

### Software Availability

*DTWax* is open-sourced here: https://github.com/harisankarsadasivan/DTWax.

